# Pathway-enhanced Transformer-based robust model for quantifying cell types of origin of cell-free transcriptome

**DOI:** 10.1101/2024.02.28.582494

**Authors:** Shuo Yan, Xuetao Tian, Yulong Qin, Jiayi Li, Tingyu Yang, Dongyue Yu, Wentao Liu, Jinghua Sun, Chengbin Hu, Qing Zhou, Zhongzhen Liu, Wen-Jing Wang

**Author notes:** These authors contributed equally: Shuo Yan and Xuetao Tian.

## Abstract

Analyzing cell types of origin of cell-free RNA can enhance the resolution of liquid biopsies, thereby deepening the understanding of molecular and cellular changes in development and disease processes. Existing deconvolution methods typically rely on meticulously curated gene expression profiles or employ deep neural network with vast and complex solution spaces that are difficult to interpret. These approaches overlook the synergistic and co-expression effects among genes in biological signaling pathways, compromising their generalizability and robustness. we developed ‘Deconformer’, a Transformer-based deconvolution model that integrates biological signaling pathways at the embedding stage, to address these issues. Compared to popular methods on multiple datasets, Deconformer demonstrates superior performance and robustness, and is capable of tracking the developmental process of the fetal and placenta. Additionally, pathway-level interpretability of Deconformer offers new insights into crosstalk, dependencies, and other interactions within cell-free RNA pathways, supporting further biological discoveries. We posit that Deconformer represents a significant advancement in the precise analysis of the cell-free transcriptome. It holds the promise of describing disease progression and severity with a new level of accuracy, focusing on the contributions of originating cell types and their pathway dependencies. This model has the potential to catalyze the development of non-invasive diagnostic tools and enhance our understanding of the underlying biology of diseases.

## 1 Introduction

Cell-free RNA (cfRNA) is a mixture of transcripts continuously released into body fluids by tissues throughout the body[1]. Previous researches have highlighted cfRNA’s potential as a powerful noninvasive measurement for health monitoring, disease prediction, diagnosis and prognosis[2–5], with its detectability in biofluids like plasma, saliva, and urine underscoring this point[6]. Analysis of the origins of cfRNA is crucial for cancer diagnosis and monitoring the development of organs and progression of diseases[2, 4], aiding in understanding diseases and development in conjunction with cellular physiology. Existing studies employing specific gene expression profiles (GEPs) for cfRNA deconvolution have traced origins to only a limited range of cell types[1, 5, 6]. Vorperian et al. [7] attempted to use CIBERSORTx[8] to generate cell-specific GEPs for deconvolution of plasma cfRNA (CSx-Vorperian), aiming to obtain relative fractions of cfRNA origins for a vast majority of cell types. However, compared to universally expressed housekeeping genes, genes with a high power to distinguish cell types tend to have lower expression levels[9] and higher missing rates, the absence and bias of these specific genes in expression profiles greatly affect the accuracy and stability of regression or factorization algorithms in deconvolution. To enhance robustness against input noise and technical biases, Menden et al. developed Scaden[10], a deep neural network (DNN) method based on multilayer perceptron (MLP) for bulk RNA deconvolution, using gene expression information to infer the cellular composition of tissues without relying on complex specific gene selection. However, the solution space of MLPs is vast and entirely data-driven, lacking an understanding of inherent correlations inside data, and optimization process is unstable. Although the hidden layers of DNNs learn higher-order latent representations of cell types, this makes interpreting the model’s workings difficult. Also, most GEP-based methods require regression calculations for each sample[8, 11], and Scaden needs separate models trained for every different expression profile[10, 12], both incurring significant time costs.

Previous deconvolution methods integrate expression profiles from multiple cell types and use genes as the smallest unit to enumerate cell types from transcript mixtures. However, one challenge of cfRNA-based liquid biopsies is its high variability, manifesting as dropout of low abundant but informative transcripts. Gene-based analyses overlook that gene interactions often exist in the form of networks. Physiological and cellular processes often affect groups of synergistically acting genes[13–15], with this synergy and co-expression effect being more pronounced in the release process of cfRNA. Some cell type annotation methods have employed pathway information to improve their performance. TOSICA[16] incorporated pathway information to enhance embedding, but its computational complexity and massive parameter quantity limited the inclusion of numerous pathways or gene sets. scBERT[17] experienced a loss in continuous value embedding when using gene co-expression information for embedding.

To address these challenges, we developed Deconformer (deconvolution Transformer), a deconvolution model based on Transformer[18] that integrates pathway information during the embedding process. Transformer has become a foundational model in many fields[19–22], allowing Deconformer to be trained on hundreds of thousands of cfRNA samples simulated using single-cell RNA sequencing (scRNA-seq) data and automatically infer cell fractions without relying on predefined GEP. By utilizing pathway prior knowledge, Deconformer achieve lossless embedding of continuous gene expression values, and the addition of these synergistic and co-expressed gene information endowed Deconformer with better representation capabilities. Moreover, the tiny and efficient characteristics of model’s embedding allows for a significant increase in the number of included pathways or gene sets, even covering all pathways within a collection. Deconformer exhibits greater robustness, noise resistance, and dropout handling than previous methods. Also, its outstanding generalization capability enables the model, once trained, to infer cfRNA expression profiles obtained by different sequencing technologies. Additionally, since Deconformer Utilizes the self-attention mechanism at the pathway level, it provides pathway-level interpretability. This offers new insights into crosstalk, dependencies, and other interactions at the cfRNA pathway level, supporting further biological discoveries. To our knowledge, Deconformer pioneers the application of transformer architecture in transcriptomic data deconvolution and innovatively integrates pathway information to design a tiny, efficient, and lossless method for embedding gene expression values.

## 2 Results

### 2.1 The Deconformer algorithm

Deconformer is a cfRNA deconvolution tool based on the Transformer encoder (Fig. 1). It learns to deduce cell type fractions through supervised training on simulated data, extracting insights from high-dimensional gene expression information by incorporating biological pathways and self-attention mechanisms.

**Fig. 1.**
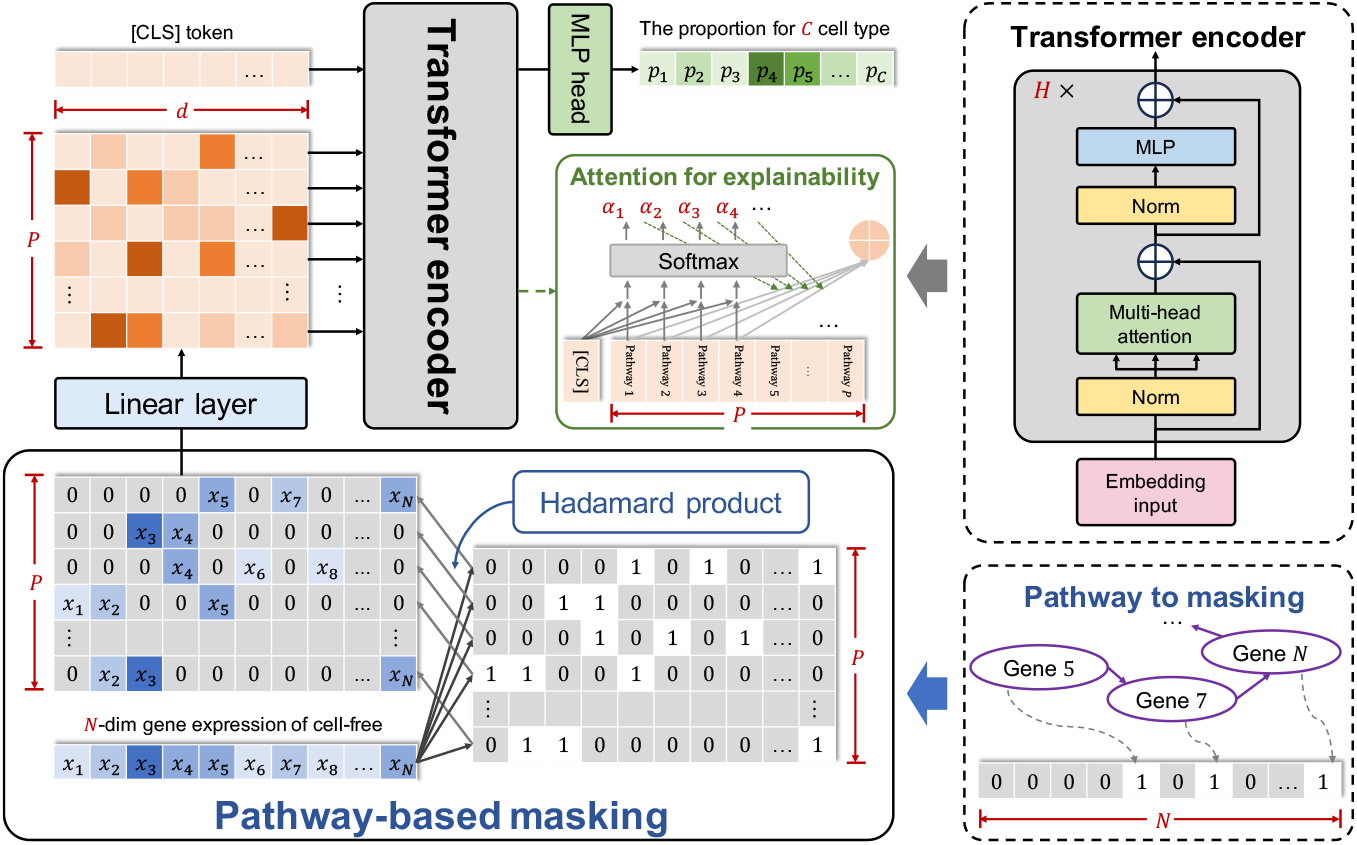
The Deconformer model overview. For each input sample’s expression profile, it is first extended via a row-wise replication to form a matrix of the same size as the mask matrix and then subjected to a Hadamard product with the mask matrix. The resulting matrix is then mapped to another space through a linear layer, and a `[CLS]` representation is added at the head. After that, multi-head self-attention is calculated, and the updated `[CLS]` representation, following a fully connected layer and softmax, yields the cell type fractions.

Deconformer consists of three parts: Pathway-based embedding, Transformer encoder, and fractions prediction. Pathway-based embedding integrates gene expression value and pathway information organically to obtain a low-dimensional representation. Firstly, a pathway-based mask matrix is constructed using pathway information covering genes with high variance in the simulated data. The cfRNA expression profile is row-wise replicated to match the size of the mask matrix, and a Hadamard product is calculated with the mask matrix to produce a masked expression matrix. Consequently, the original expression vector is dimensionally expanded into a matrix that represents expression across various pathways. Then, to capture effective information between pathways, each row of the expression matrix are mapped to the same low-dimensional space via linear transformation. The generated matrix is combined with an extra `[CLS]` representation[19] at header and further input into Transformer. After processing through each layer of the self-attention mechanism, the `[CLS]` representation integrates information from all pathway corresponding representations and is then transformed into a vector for predicting cell type fractions through a fully connected network. Biological processes are complex and interactive and Deconformer’s biologically meaningful embedding allows the self-attention mechanism to obtain the attention scores between pathways, indicating the interactions between pathway pairs, and that over `[CLS]` representation, reflecting each pathway’s contribution. Furthermore, the capability of including all pathways within a single pathway dataset and abandoning the use of dimensionality reduction enables Deconformer to keep the full pathway-level interpretation.

### 2.2 Performance on simulated cfRNA and distinct body fluid cfRNA dataset

We compared Deconformer with the previously reported cfRNA deconvolution method CSx-Vorperian and Scaden, known for its superior performance on bulk data. CSx-Vorperian relies on manually curated GEP matrices but can also use single-cell data to generate corresponding GEP matrices. The Scaden method, as described in the original paper, was trained using simulated data consistent with Deconformer. We separately simulated 1000 cfRNA samples for performance testing. The results showed that Deconformer achieved the highest Lin’s concordance correlation coefficient (CCC)[23], Pearson’s correlation coefficient (*r*), and the lowest root mean square error (RMSE) on simulated data (Fig. 2A). Given the non-negligible impact of dropout in cfRNA sequencing[24], we further tested the robustness of the method by evaluating the inference performance of the three methods under different dropouts. This was achieved by randomly dropping expression in these 1,000 simulated cfRNA profiles at rates of 0.1, 0.2, 0.3, 0.4, and 0.5. Deconformer still performed the best on the aforementioned evaluation metrics. Moreover, under different dropout rates, the performance degradation (loss) and its extent of Deconformer were smaller than that of DNN-based Scaden and regression-based CSx-Vorperian(Fig. 2B, 2C and Extended Data Fig.1A). Notably, on simulated data, Deconformer’s CCC reached over 99%, and even with a 50% dropout rate, the CCC remained above 95%.

**Fig. 2.**
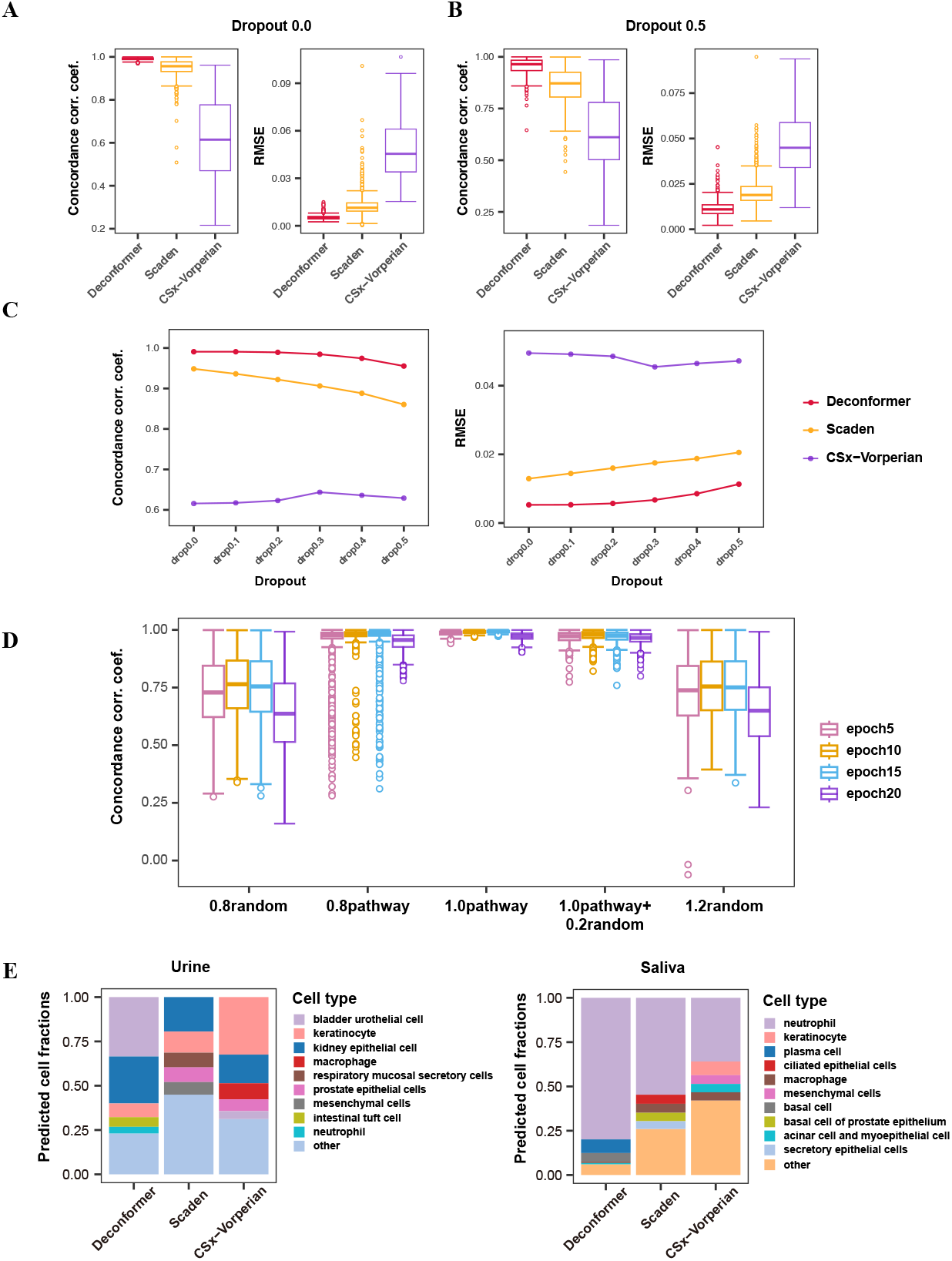
Evaluation of accuracy and generalization ability on simulated cfRNA data, as well as the importance of pathway information. A: Deconvolution performance of Deconformer, Scaden, and CSx-Vorperian on simulated data. B: Deconvolution performance of the three methods after dropping 50% of the simulated data expression profile. C: Deconvolution performance of the three methods on simulated data under different dropout conditions. D: Performance of Deconformer models trained with different perturbed and completely random mask matrices on simulated data. E: Top-5 cell types and the sum of ‘other’ for different body fluids, urine (left), and saliva (right), across the three methods.

To assess the significance of pathway information to Deconformer, we performed an test involving progressive perturbations on the original pathway mask matrix, including a 20% reduction of non-zero values to simulate partial information loss, and a 20% increase of random ones to mimic noise. Additionally, we generated two completely random mask matrices with non-zero element ratios matching the perturbed matrices for comparison. Train deconformer using these mask matrices separately. The results indicate that, whether with the original expression profiles or with 0.5 dropout, reducing non-zero values by 20% achieved the best performance on CCC and RMSE, followed by adding 20% noise, yet still not as effective as the original mask matrix. The completely random mask matrices consistently had CCC values below 0.8 across different training epochs, indicating poor performance that could not be improved by increasing training epoch (Fig. 2D and Extended Data Fig.1B,1C). This demonstrates that the potential co-expression information in the pathway mask can effectively enhance deconvolution performance, and the self-attention mechanism can better capture the true correlations between pathways.

To test Deconformer’s generalization performance on cfRNA expression profiles generated from different body fluids, we conducted deconvolution on urine, saliva, seminal plasma, and blood plasma[17]. The results showed that Deconformer could identify the patterns of different body fluids, and the top five contributing cell types for each fluid source were more consistent with expectations compared to Scaden and CSx-Vorperian. For example, the primary sources of cfRNA in urine were the bladder and kidneys (Fig. 2E and Extended Data Fig.1D).

### 2.3 Accurate identification of cancer lesion differential signals

cfRNA shows high specificity in early cancer screening, presenting promising prospects[5, 25–27]. Differentiating transcripts of lesion-specific cells from high-background transcripts originating from blood cells is crucial for early cancer screening and detection. We performed tests on hepatocellular carcinoma (HCC), colorectal cancer (CRC), gastric cancer (STAD), multiple myeloma (MM), and their precancerous conditions to demonstrate that Deconformer accurately and stably identifies changes in cfRNA origin fractions from tissues and organs directly affected by the disease or involved in both solid and non-solid tumors. Generally, lesions affect the surrounding tissues and cell types, as well as the immune system, The affected cell types include those with definite changes in fraction scores or associated cell counts, as well as those anticipated to potentially undergo changes in source fractions. To emphasize the advantages and potential limitations of Deconformer, we compared its conclusive results only with CSx-Vorperian and Scaden.

We first conducted tests on solid tumors. Hepatocyte damage is a prominent feature of HCC, we deconvolved samples from the Chen et al. [25]and Roskams-Hieter et al. [26] datasets using the aforementioned three methods. The results of deconvolution demonstrate a significant increase in the hepatocyte fractions in the patient group across both datasets, a difference not detected by Scaden in the Roskams-Hieter et al. dataset, and although CSx-Vorperian showed a difference. While CSx-Vorperian identified a difference, the control group primarily consisted of individuals with zero hepatocyte fractions.(Fig. 1A and Extended Data Fig.2A). Through the deconvolution of datasets from Chen et al. and Tao et al. [27], Deconformer observed an increase in platelet fractions in both CRC groups, which aligns with the common thrombocytosis observed in patients with CRC. Additionally, Deconformer discovered a decrease in the fractions of cell types that are closely related to the lifecycle and play a crucial role in the self-renewal processes of the intestine, such as immature intestinal epithelial cells which have been proven to be directly related to c CRC[28, 29].(Extended Data Fig.2B). In the case of gastric cancer, both datasets demonstrated a decrease in the contribution scores of stromal cells, NK cells, monocytes, B cells, neutrophils, and duodenum glandular cells[30],(Extended Data Fig.3A), showcasing the stability of Deconformer across datasets. To further validate the cell pathophysiological significance and disease discrimination ability of cell type fractions, we used the deconvolution results from Tao et al.’s data for classification modeling of two types of cancer. The results showed a performance comparable to the best multi-omics results in CRC and better than cfRNA abundance models in STAD, possibly due to the reference cell types included.(Extended Data Fig.3B)

In non-solid tumors, employing Deconformer on MM data from Roskams-Hieter et al. dataset, we observed a significant increase in plasma cells and a marked rise in erythrocyte and erythroid progenitor in monoclonal gammopathy of undetermined significance (MGUS) group. In the MM group, there was not observe a significant increase in plasma cells, but we also noticed an increase in erythrocyte and erythroid progenitor (Fig. 3B and Extended Data Fig.3C). MM and MUGS generally present with anemia, but our observation of an increase could be due to the uncontrolled abnormal proliferation of plasma cells and cancerous plasma cells in the bone marrow, compressing normal erythrocyte and erythroid progenitor, causing damage and apoptosis of these cells, leading to anemia and a corresponding increase in cfRNA signals in plasma. The increase in plasma cells in MUGS can well explain the disease’s onset and progression. The absence of increased plasma cells in the MM led us to speculate significant changes in RNA expression between cancerous and normal plasma cells in MM patients[31], causing Deconformer to fail to recognize cancerous plasma cells.

**Fig. 3.**
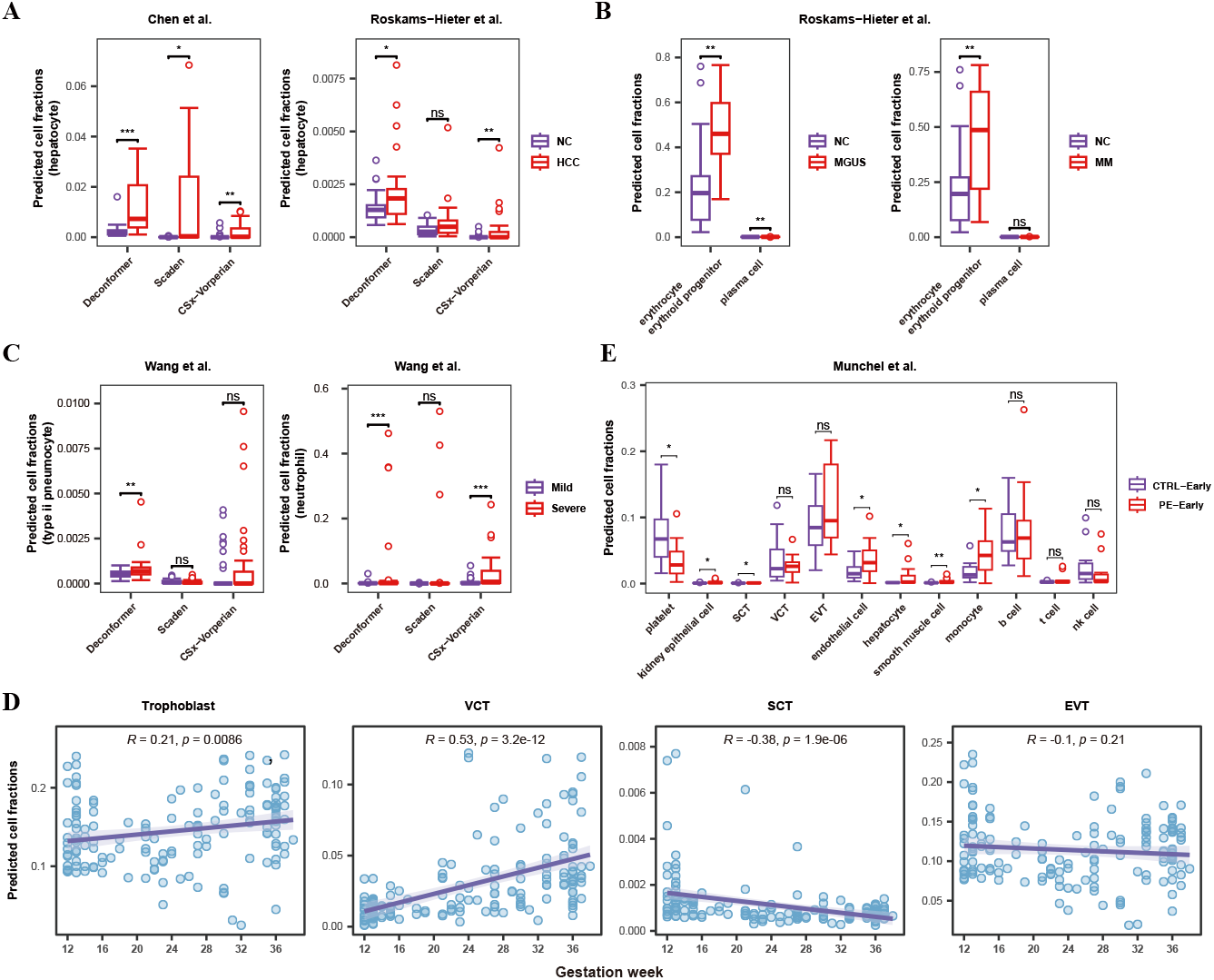
Comparison of the ability to differentiate cell types on disease differences and Deconformer can track organ development. A: Capture of liver signal differences between liver cancer and healthy groups by Deconformer, Scaden, and CSx-Vorperian in the two HCC datasets of Chen et al. and Roskams-Hieter et al. B: Differences in cell type fractions scores in MGUS and MM as captured by Deconformer. C: Differences in type ii pneumocyte and neutrophils between severe and mild COVID-19 patients by Deconformer, Scaden, and CSx-Vorperian in Wang et al. data. D: The changes in fractions of trophoblast and VCT, EVT, SCT over gestational weeks. E: Differences in cell type fractions between early-onset preeclampsia and healthy individuals in Munchel et al. pregnancy data as captured by Deconformer.

### 2.4 Changes in affected organs and peripheral immunity due to viral infection

To further demonstrate Deconformer’s ability to capture differences in disease progression at lesion sites and in peripheral immunity for viral infection diseases, we applied Deconformer to datasets on coronavirus disease 2019 (COVID-19) infection by Wang et al. [32]and Hepatitis B virus (HBV) infection by Sun et al. [33]. COVID-19, an infectious disease caused by the Severe Acute Respiratory Syndrome Coronavirus 2 (SARS-CoV-2), primarily exhibits diffuse alveolar damage in severe cases[34]. Deconformer’s deconvolution results indicated a higher fractions of type ii pneumocyte cells in severely ill patients compared to those with mild symptoms, a distinction not apparent in results from Scaden and CSx-Vorperian. Regarding changes in peripheral immunity, the referenced dataset[32] showed significantly higher neutrophil counts in severely ill COVID-19 patients than in those with mild symptoms, a finding consistent with both Deconformer and CSx-Vorperian results, whereas Scaden failed to detect any differences in neutrophils (Fig. 2C). Additionally, we observed an increase in monocytes[35] in severe cases in Deconformer’s results, corroborating previous studies. The rise in T cell and NK cells, contrary to the previously reported decline in circulating T and NK cells in severe COVID-19[36–39], suggests that the depletion of immune cell subgroups may release more signals into the plasma. Besides these immune cells, we also noted a decrease in platelets in severely ill patients, aligning with the differences in blood cell counts reported in the corresponding studies.(Extended Data Fig.4A). By deconvoluting the sample data from the study by Sun et al. [33] we observed the progressive changes in hepatocyte fractions across different serological phenotypes of HBV infection. Compared to normal individuals, patients with the negative HBeAg serological pattern showed increased hepatocyte fractions,(Extended Data Fig.4B), but this increase was even more pronounced and significant in patients with the positive HBeAg pattern[33]. Additionally, in the positive HBeAg patient group, we observed a decrease in the fractions of NK cells and T cells. Existing research has illustrated that these differences could be due to a reduction in NK cell fractions[40] and T cell exhaustion[41, 42],(Extended Data Fig.4C) reflecting more severe liver damage and immune characteristics in patients with chronic infection, particularly those with the positive HBeAg serological pattern. The above differences indicate that Deconformer can infer changes in peripheral immunity for acute and chronic viral infections of different courses and severities from cfRNA expression profiles.

### 2.5 Detecting and tracking fetal organ and placenta development

The deconvolution of cfRNA offers a comprehensive means to monitor maternal and fetal organ health and to track fetal development. Moufarrej et al. [43] qualitatively observed changes in the mother and placenta during fetal development through cell and tissue-specific scores, but these results do not reflect the relative developmental changes between different organs of the individual. To track the development of fetal organs and changes during maternal pregnancy, we added fetal single-cell reference data and placental single-cell data into the reference dataset, and re-simulated cfRNA, re-training Deconformer to obtain a model specific for pregnancy stages. Before deconvoluting real cfRNA data, we generated cfRNA samples consisting only single cell type to avoid interference between maternal and fetal origin fractions, used to evaluate whether Deconformer can distinguish between maternal and fetal origin of cfRNA. The results show that Deconformer can effectively distinguish the origin, with over a thousand-fold difference in origin fractions between maternal and fetal cells of the same type, demonstrating Deconformer’s strong discriminative power for cell types with similar expressions.(Extended Data Fig.5A,5B)

Using pregnancy stages Deconformer model to deconvolve healthy pregnant women’s plasma cfRNA profile[44], We observed that the fractions of villous cytotrophoblast (VCT) most notably increases with gestation, extravillous trophoblast (EVT) does not show significant changes, syncytiotrophoblast (SCT) is on a downward trend. The overall fractions of the trophoblast (the sum of SCT, VCT, EVT) increase with gestation (Fig. 3D). Previous research[45] has shown that the shell of VCT gradually diminishes and becomes sparse, and that VCT cells remain in a proliferative state throughout pregnancy. The primary function of the EVT is invasive remodeling, which increases in the early stages of pregnancy and eventually reaches a state of equilibrium. Meanwhile, the increased proportion of other cell types may lead to a passive reduction in the fractions of SCT. All these possible scenarios are consistent with the results from the Deconformer model. Among fetal cells, the fractions of mesencephalon and hepatocyte cells increased significantly with gestation, while cardiomyocyte cells significantly decreased, and there was no significant change in the fractions of kidney epithelial cells. In the early-onset pre-eclampsia (PE) group, Deconformer showed significantly higher contributions of endothelial cell, kidney epithelial cell, smooth muscle cell, hepatocyte, monocyte[46] compared to the control group, consistent with the symptoms of proteinuria, impaired hepatocyte function, renal insufficiency, and increased blood pressure in early-onset PE patients[43] (Fig. 3E), and an increased contribution of platelets, possibly related to thrombocytopenia in early-onset PE patients. But these changes and differences were not observed in the late-onset PE group, which also corroborates, from the changes in affected organs, that early-onset PE is more severe than late-onset PE. It is noteworthy that in both early-onset and late-onset PE, the SCT fractions significantly increased, even though the absolute values of the SCT scores are very small.(Extended Data Fig.5C). Additionally, For Intrahepatic Cholestasis of Pregnancy (ICP) patients in the Sun et al. [33] dataset, using pregnancy stages Deconformer model, we found a significant increase in the hepatocyte fractions, consistent with reports. The Deconformer with adult reference cells yielded the same result, demonstrating Deconformer’s robustness.(Extended Data Fig.5D)

### 2.6 Deconformer can identify dependencies between pathways in cfRNA

Previous deconvolution methods struggled to adequately explain or provide insights into differences in cell type origin fractions. We obtained the last layer’s attention scores from the inference stage of samples from COVID-19 and HCC datasets. Our interpretability analysis is in two parts: First, we analyzed the attention scores between the `[CLS]` representation used for calculating cell fractions scores and all pathways. We calculated the Cohen’s *d* effect of `[CLS]` representation attention scores for different groups, using pathways with an absolute value greater than 0.8 for self-clustering (Fig. 4C). On both datasets, this method demonstrates better discriminating ability for samples of different groups than differential genes selected with the same Cohen’s d threshold, reflecting Deconformer’s noise reduction capability to a certain extent. Specifically, for the neutrophil activation-related pathways enriched differently between mild and severe cases as discovered by Wang et al., the attention score of the `[CLS]` representation also exhibits significant differences.(Extended Data Fig.6A)

**Fig. 4.**
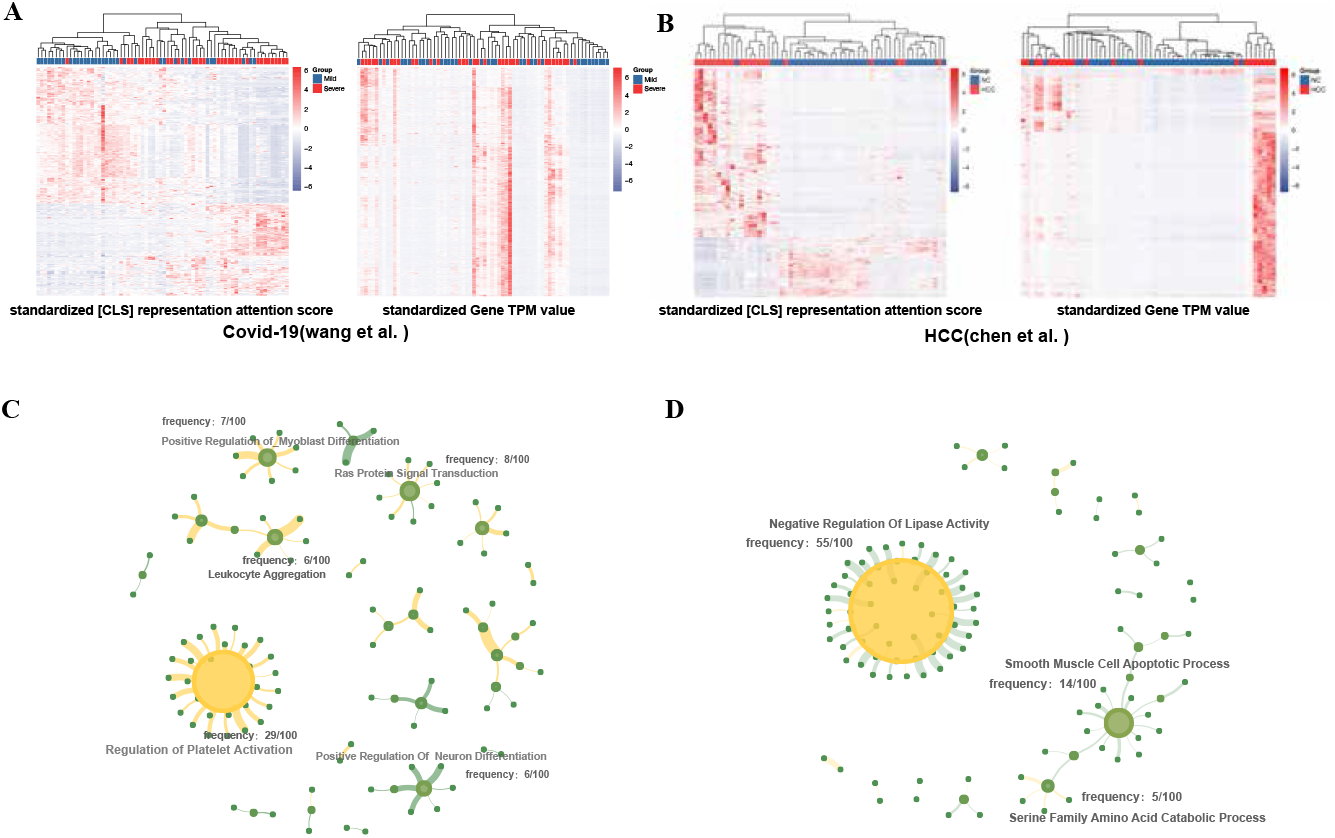
The dependency relationships of pathways in plasma cfRNA have biological significance. A: Heatmaps of unsupervised clustering of significant differences in attention scores and gene expression profiles in COVID-19 data from Wang et al. B: Heatmaps of unsupervised clustering of significant differences in attention scores and gene expression profiles in HCC data from Chen et al. C: Interactions and dependencies between pathways in plasma cfRNA of COVID-19 patients as identified by attention score between all pathway pairs. D: Interactions and dependencies between pathways in plasma cfRNA of HCC patients as identified by attention score between all pathway pairs.

To further explore interactions between pathways in plasma cfRNA, we considered the overall correlations and dependencies among 5,000 pathways. We performed Cohen’s *d* effect statistics on 25 million pathway pairs between case and control groups, selecting the top 100 pairs with the highest absolute Cohen’s *d* effect for pathway network analysis. We found that Regulation of Platelet Activation, which appeared most frequently among these differential pathway pairs, might influence many physiological processes related to severe COVID-19, consistent with previous reports[47–49]. Additionally, pathways such as Leukocyte Aggregation[50], Ras protein signal Transduction, and Positive Regulation of Myoblast Differentiation, which have been proven related to severe COVID-19, also showed high differences (Fig. 4D). In the HCC data from chen et al., we found pathways like Negative Regulation of Lipase Activation[51], Smooth Muscle Cell Apoptotic Process, and Serine Family Amino Acid Catabolic Process[52] showed the most dependencies with other pathways (Fig. 4E). similar results were observed for both types of analysis on HCC data from Roskams-Hieter et al.(Extended Data Fig.6B,6C).

These findings indicate the self-attention mechanism effectively learns the interactions between pathways. Additionally, for pathways which differences were not detected in the enriched expression profiles, Deconformer can still effectively identify potential key pathways, offering a new perspective on the interactions between biological pathways in plasma cfRNA for diseases and discovering important and central pathways.

## 3 Discussion

To address the robustness and accuracy of cfRNA deconvolution and the interpretability of critical pathways, we developed Deconformer. It learns patterns between pathway interactions and cell type fractions in simulated cfRNA gene expression profiles for inferring cell type fractions in real cfRNA expression spectra. Deconformer is the first to apply the transformer to transcriptomic data deconvolution. It explicitly incorporates prior knowledge into the model’s embedding module, using gene groups with synergistic and co-expression effects as the smallest unit for modeling, rather than individual genes susceptible to random noise. Moreover, its embedding design is lightweight and efficient, accommodating nearly ten thousand pathways or gene sets, enhancing the model’s comprehensive understanding of inter-pathway interactions.

Deconformer demonstrates a particularly notable robustness to noise and bias. With a single training session, it can infer expression profiles obtained from different bodily fluids and sequencing technologies, surpassing existing advanced methods in both simulated and multiple real datasets. If only using pre-trained model parameters for inference, a laptop without GPU is sufficient, eliminating the need for training from scratch. Additionally, Deconformer can flexibly track placental and fetal development by adjusting single-cell reference datasets.

We hope Deconformer advances the understanding of pathway-pathway interactions in cfRNA from plasma, directly modeling small sample data, which is difficult for complex models capturing pathway interactions in bodily fluid cfRNA. Through Deconformer’s deconvolution process, this objective is also achieved. In COVID-19 data, using Deconformer’s attention scores, we mapped the dependency network between pathways. The most significant differences in pathway-pair attention levels and network hub pathways were in platelet activation, directly related to severe cases. This offers important pathways for studying disease mechanisms and treatments in the peripheral immune system and supports further biological discoveries. Concurrently, this will help deepen our definition and understanding of the role of cfRNA.

Although Deconformer achieved good performance on the samples and data used in this study, it is important to note that cell type similarities, sample heterogeneity and complexity, and experimental noise and bias significantly limit deconvolution accuracy. Interpretation of individual results should be approached with caution. In the future, as single-cell datasets become more comprehensive, more detailed cell types like immune cells residing in different tissues and cancer cells can be added to the reference data to obtain relative fractions of circulating and resident immune and cancer cells. Currently, Deconformer’s pathway-related genes consist of mRNA. However, non-coding RNAs are also crucial in disease research, and integrating functional pathways of non-coding RNAs into Deconformer is equally important. Also, grouping pathways by function and adding positional encoding to tokens might enhance the model’s representational power and training speed. Of course, Deconformer’s structure can be easily adapted to other tasks, and we hope to explore its flexibility and versatility in various downstream tasks, such as cell type annotation and disease differential studies. By expanding the cell types and distributions in simulated data, the future Deconformer is expected to become a general model for whole transcriptome data deconvolution and annotation.

## 4 Methods

### 4.1 Single-cell datasets and preprocessing

The single-cell reference dataset for training the adult model simulation data was sourced from the Tabula Sapiens project (TSP), containing approximately 500,000 single cells across 175 cell types. Considering the similarity in expressions among cell types, we followed the cell type merging method of Vorperian et al. for the TSP cell types. Moreover, we retained only cell types with more than 10 cells to ensure diversity in the single-cell data selected for simulation accurately represents each cell type’s characteristics, resulting in a total of 60 merged adult cell types. (Supplementary Notes 1)

For simulating pregnancy stage model training single-cell data, in addition to TSP, we included a placental single-cell dataset obtained from Vento-Tormo et al. and fetal single-cell data from Xu et al. Taking detectability and batch effect interference into account, only the most representative cell types were chosen from these three sources. The placental single-cell data comprised placental trophoblasts, including SCT, EVT, VCT, while fetal single-cell data covered representative cell types from the four main fetal organs: heart, liver, brain, and kidney. For maternal cell types, we preserved the top-25 cell types identified by Deconformer model inference from the pregnant women’s dataset, along with the cell types covered by fetal cell types. In summary, the comprehensive collection consisted of 4 fetal cell types, 3 placental trophoblast types, and 27 adult cell types, totaling 34 cell types.(Supplementary Notes 1)

Only cells from droplet sequencing (’10x’) were used for analysis in these datasets. For the TSP dataset, raw count values from the DecontX adjusted layer were used to minimize signal spread contamination in individual cells. The expression values of individual cells were normalized to 10,000 using the “normalize total” function from scanpy. Since the genes in the pathways used are predominantly mRNA, only mRNA was retained for subsequent analysis. These values were used as input vectors for generating simulated cfRNA.

### 4.2 Simulated cfRNA data

Deconformer was trained on simulated cfRNA data, generated as follows: 1. Randomly sample 2 to *S* cell types from a pool of total original types with equal probability. 2. For each cell type, randomly sample 200 to 800 cells for the mixture. If the cell number of a particular type is less than 200, all available cells were included in the mixture. 3. Specify the mixture fractions for the cell types by assigning a random ratio to each cell type with the sum of all ratios being 1. 4. Accumulate the expression data for the cell types, where the expression value of each cell type was first averaged and then aggregated into a weighted sum based on the mixture fractions. Based on these phases, we simulated 800,000 cfRNA samples, as well as an additional 1,000 cfRNA samples for performance testing. In this study, *S* was set as 20 (1/3 of total cell types) for adult model and 17 (1/2 of total) for pregnancy stage model,with no need for users to set other parameters.

### 4.3 The proposed Deconformer model

Gene abundance of cell-free transcriptome is the accumulation of genes from all origin cells and can be represented as a *N* -dimensionality vector 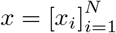, where *N* denotes the number of genes and *x*_*i*_ denotes the expression value of *i*-th gene. Taking the cfRNA expression profile as input, a deconvolution model will output the fractions of each type of cell to all cells, represented by a *C*-dimensionality vector 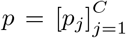, where *C* is the number of cell types and *p*_*j*_ is the fractions of *j*-th type of cell. In this article, we design a novel Deconformer model, which leverages prior knowledge of pathways, adopts the advanced paradigm of Transformer and tailors its architecture specifically for the task of cfRNA deconvolution. On one hand, pathway information indicates which genes have certain associations and is beneficial for the model to solve faster and more accurately. On the other hand, Transformer has a global receptive field, enabling it to effectively learn global representations and long-range dependencies, and achieves excellent performance on non-sequential data. The details of Deconformer model are presented from the following three aspects:

#### 4.3.1 Pathway selection

The pathways in this study were constructed using the C5 GOBP dataset (Gene sets derived from the GO Biological Process ontology) from the GSEA database. Each pathway in C5 GOBP consists of several genes and thus we evaluated its value by whether it covers more useful genes. Specifically, the top-*G* genes having the highest between-subject variance are selected and used as evaluation criteria. The Jaccard similarity between each pathway and the *G* genes is calculated, and the top-*P* pathways having the biggest similarity are selected for further modeling. In this study, *G* and *P* are set as 10, 000 and 5, 000 respectively.

#### 4.3.2 Pathway-based embedding

As above-mentioned, the input of the model is *N*-dimensionality gene expression 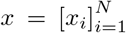. To model its different representations over each pathway, *x* is first extended via a row-wise replication and the obtained matrix is denoted as *M*_*X*_ ∈ ℝ^*P ×N*^. Meanwhile, according to the involved genes of the selected *P* pathways, a pathway-based mask matrix is established, denoted as *M*_*P*_ ∈ {0, 1}^*P ×N*^, where each element is valued as 0 or 1, representing whether a pathway covers one specific gene. As Equation (1), the mask matrix *M*_*P*_ act on *M*_*X*_ with Hadamard product to achieve a new representations *M*_*E*_ ∈ ℝ^*P ×N*^.

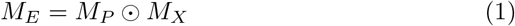

Then, the matrix *M*_*E*_ is transformed into a deeper *d*-dimensionality space through a linear layer as Equation (2), where *W*^*E*^ ∈ ℝ^*N ×d*^ is the trainable parameters and *M*_*T*_ ∈ ℝ^*P ×d*^ can be seen as the representations of the input gene expression over different pathways. In this study, *d* is set as 128.

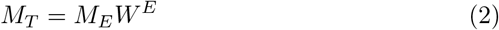

#### 4.3.3 Model building

On the basis, Transformer module is leveraged to capture the intrinsic correlation among the representations over different pathways. Concretely, an extra trainable *d*-dimensionality `[CLS]` representation is randomly initialized using a normal distribution and added to the header of *M*_*T*_, resulting in a representation list *T* ∈ ℝ ^(*P* +1)*×d*^ as the input of Transformer. The representation list *T* is encoded by four consecutive layers of Transformer, where the original positional embedding of Transformer is not utilized because there is no positional relationship between the pathways. Each layer of Transformer consists of multi-head attention layer and the following residual connection, layer normalization, as well as feedforward neural network layers. The process with input representation list being *X* ∈ ℝ ^(*P* +1)*×d*^ is implemented by the following steps.

a. **Multi-head attention**. The layer performs *H*-time self-attention operations. The *h*-th attention is performed as Equation (3), (4) and (5). In this study, *H* is set as 4.

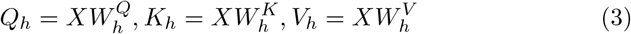

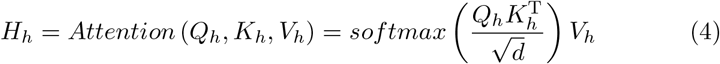

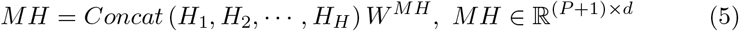

where 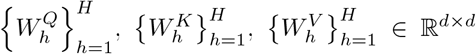 and *W*^*MH*^ ∈ ℝ^*Hd×d*^ are trainable parameters.
b. **Residual connection and layer normalization**. This layer adds the original input *X* and the multi-head attention output *MH* together, and applies layer normalization over this as Equation (6).

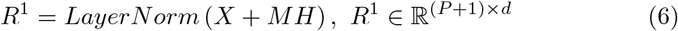
c. **Feedforward neural network**. In this layer, each representation 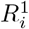 in *R*^1^ is updated by a two-layer neural network as Equation (7).

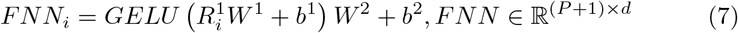

where 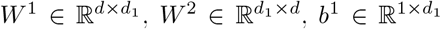 and *b*^2^ ∈ ℝ^1*×d*^ are trainable parameters, and *GELU* (•) is the activation function with Gaussian error linear unit. In this study, *d*_1_ is set as 512.
d. **Residual connection and layer normalization again**. This layer adds *R*^1^ and *FNN* together, and execute layer normalization again as Equation (8). Finally, one Transformer layer outputs a representation list of the same size as the input, i.e. *R*^2^.

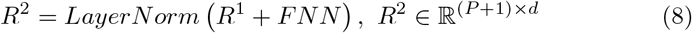

Through these four Transformer layers, the representation list *T* is transformed into 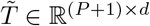, which embeds rich correlation information between pathways. Especially, the corresponding representation at the `[CLS]` position, 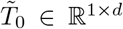, is utilized to further predict the fractions of each cell type as Equation (9).

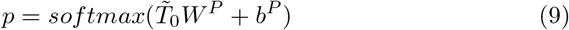

where *W*^*P*^ ∈ ℝ^*d×c*^ and *b*^*P*^ ∈ ℝ^1*×c*^ are trainable parameters. Each element of 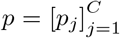 represents the fractions of corresponding cell type, and their sum is 1.0.

#### 4.3.4 Model training

Due to the inevitable noise and biases in the expression profiles obtained from cfRNA sequencing, to encourage the model to learn more complex representations of cell types, 20% of input values are randomly set to zero before the simulated cfRNA data is inputted into the model. The proposed Deconformer model is optimized using a mean squared error (MSE) loss between the prediction *p* and ground-truth 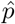 as Equation (10), where *I*( •) is indicator function, and train it by mini-batch Adam optimizer with an end-to-end strategy. Training hyperparameters were tuned to the following values: max learning rate, 0.0005; learning scheduler, linear with warmup; warmup steps, 10000; batch size, 128. Typically, Deconformer converges within 15 epochs. Training for more epochs does not significantly increase the CCC on simulated data and poses a risk of overfitting. (Fig. 2D)

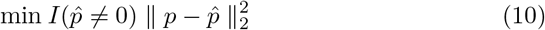

### 4.4 Model interpretability

The Deconformer model is based on a self-attention mechanism, and we conducted interpretability analysis to explore key pathways in decision-making. The attention score reflect the contribution of each pathway and the impact of interactions between pathways on the outcome. Since our final output is a multi-dimensional vector, and certain cell types affected by diseases have relatively low origin fractions, we did not directly use the magnitude of self-attention scores. Instead, we utilized pathways with the highest inter-group differences. These pathways with the greatest differences reflect the pathways that have the largest impact on score variances under different disease states. Additionally, these pathway pairs, forming a differential network, exhibit potential interactions and dependencies.

Attention score can be obtained from Equation (4), and we averaged the attention scores from multiple heads, integrating attention matrices from multiple heads into one matrix. In this average attention matrix, each value represents the degree of attention from pathway *i* to pathway *j*. We first analyzed the attention scores between the `[CLS]` representation and other pathway representations, which is a vector of the length of the number of pathways. This way, we can obtain the degree of attention of the `[CLS]` representation to other pathways in different states of samples used for classification. Secondly, to obtain differences in attention degrees between each pathway, we also analyzed the attention between 5000 pathways, i.e., 25 million pathway pairs. The calculation of differences in pathways used the Cohen’s *d* statistic, and the pathways were ranked by the absolute value of their differences.

### 4.5 Comparison methods

For benchmarking, we selected two representative tools, CSx-Vorperian and Scaden, which are regression-based algorithm on GEP and based on DNN, respectively. In real data comparison, we only conducted comparisons on data with clear evidence support or corresponding blood cell counts.

#### CSx-Vorperian

CSx-Vorperian was tested following its provided instructions and tutorials. Due to its high time complexity, each sample requires at least 14 hours. For simulated data comparison, we randomly selected 100 out of 1000 validation data for performance testing. The GEP provided by Vorperian et al. includes 62 cell types and also contains lncRNA. We used the original GEP provided in the article for performance comparison with real data. Since the simulated data contains only 60 cell types and mRNA, which does not completely match the GEP provided in the article, we generated the corresponding GEP based on the method from the original article for performance comparison with the simulated data.

#### Scaden

Given the significantly larger variety of single-cell sources in the simulated data compared to those tested in the Scaden paper and the high dropout in cfRNA, the Scaden model was unable to filter out enough genes with a variance greater than 0.1 as in the original paper. Therefore, based on the instructions and source code from the original Scaden paper, we rewrote a version of Scaden using PyTorch2.0. The only change we made was in the gene selection part, where we selected the top 10,000 genes with the highest variance instead of those with a variance greater than 0.1. We then tested the impact of different training set sizes on the performance of Scaden on a validation machine and ultimately determined that performance does not improve further with training sets larger than 9,000. For performance comparison using Scaden, we used a model trained on a training set based on 9,000.

### 4.6 Performance evaluation and statistical analysis

The performance evaluation metrics for the algorithm include CCC, *r* (Pearson’s correlation coefficient) and RMSE. Relying solely on a single metric can make it challenging to evaluate performance reasonably in all scenarios. The CCC value is prone to distortion when the cell types in the expression spectrum are few. Given that the loss functions for the Deconformer and Scaden models are sparse MSE (Mean Squared Error) and MAE (Mean Absolute Error) respectively, the RMSE can also be low when the model is overfitting. Therefore, it is necessary to use both CCC and RMSE for performance assessment. Statistical differences are examined using the Wilcoxon rank-sum (Mann-Whitney-U) test.

### 4.7 cfRNA dataset and processing

The datasets for cancer-related diseases are from Chen et al., Roskams-Hieter et al., Tao et al. For infection-related diseases, COVID-19 research data is from Wang et al., HBV infection data from Sun et al. Placental development and pregnancy-related diseases are from Munchel et al., Sun et al. Bodily fluid data is from Liu et al. These data were processed to TPM, with the quality control criteria for the samples based on the rules outlined in the original articles.

### 4.8 Software and hardware environment

Deconformer utilizes Python 3.10.12 and is primarily built using torch (version 2.0.0+cu117) and scanpy (version 1.9.3). This study used the Nvidia A100 40GB to train models. If only using pre-trained model parameters for inference, a laptop without GPU is sufficient. Downstream analysis and plotting employed joblib (version 1.3.2), as well as R 4.2.0, ggplot2 (3.4.4), pheatmap (1.0.12), igraph (1.5.1), ggraph (2.1.0), and their dependent packages.

## 5 Data Availability

All data used in the article are given in the method part, and these data are publicly accessible, except for Liu et al., which was obtained from the author. Samples from Chen et al. (GSE174302), Toden et al. (GSE186607), Roskams-Hieter et al. (GSE182824) were downloaded from the Sequence Read Archive with the respective accession numbers. samples from Munchel et al. were downloaded from the Resources of the article. Samples from Wang et al. (CNP0001306) were downloaded from CNGB Sequence Archive (CNSA; https://db.cngb.org/cnsa/) of the China National GeneBank DataBase (CNGBdb) under accession number CNP0001306. Samples from Sun et al. were downloaded from Genome Sequence Archive for Human Database with accession numbers HRA003387. The code of cfRNA alignment and quantification is available at https://github.com/wonderful1/PALM-Seq-cfRNA. Tabula Sapiens was downloaded from the Chan Zuckerberg Biohub (https://tabula-sapiens-portal.ds.czbiohub.org, version 5.0). Placental single-cell data (Vento-Tormo et al.) were downloaded from ArrayExpress with experiment codes E-MTAB-6701. Fetal single-cell data (Xu et al.) were downloaded from the Gene Expression Omnibus (GEO) under accession code GSE157329.

## 6 Code Availability

The code for data preprocessing, Deconformer modeling, and downstream analysis, as well as the PyTorch2.0 version of Scaden, are available on https://github.com/findys/Deconformer. Detailed instructions are provided there.

### 7 Acknowledgement

This work was supported by grants from National Natural Science Foundation of China (62207002), and Science, Technology and Innovation Commission of Shenzhen Municipality (JCYJ20170412152854656).

## 8 Conflicts of Interest

The authors declare no conflict of interest.

## Notes

### Competing Interest Statement

The authors have declared no competing interest.

### Summary of Updates

Figure 1 revised; Figure 4 revised;

## References

[1] Koh, W. et al. Noninvasive in vivo monitoring of tissue-specific global gene expression in humans. Proceedings of the National Academy of Sciences 111 (20), 7361–7366 (2014). 10.1073/pnas.1405528111.

[2] Ngo, T. T. M. et al. Noninvasive blood tests for fetal development predict gestational age and preterm delivery. Science 360 (6393), 1133–1136 (2018). 10.1126/science.aar3819.

[3] Ibarra, A. et al. Non-invasive characterization of human bone marrow stimulation and reconstitution by cell-free messenger RNA sequencing. Nature Communications 11 (1), 400 (2020). 10.1038/s41467-019-14253-4.

[4] Toden, S. et al. Noninvasive characterization of Alzheimer’s disease by circulating, cell-free messenger RNA next-generation sequencing. Science Advances 6 (50), eabb1654 (2020). 10.1126/sciadv.abb1654.

[5] Larson, M. H. et al. A comprehensive characterization of the cell-free transcriptome reveals tissue- and subtype-specific biomarkers for cancer detection. Nature Communications 12 (1), 2357 (2021). 10.1038/s41467-021-22444-1.

[6] Hulstaert, E. et al. Charting Extracellular Transcriptomes in The Human Biofluid RNA Atlas. Cell Reports 33 (13), 108552 (2020). 10.1016/j.celrep.2020.108552.

[7] Vorperian, S. K. et al. Cell types of origin of the cell-free transcriptome. Nature Biotechnology 40 (6), 855–861 (2022). 10.1038/s41587-021-01188-9.

[8] Newman, A. M. et al. Determining cell type abundance and expression from bulk tissues with digital cytometry. Nature Biotechnology 37 (7), 773–782 (2019). 10.1038/s41587-019-0114-2.

[9] Theodoris, C. V. et al. Transfer learning enables predictions in network biology. Nature 618 (7965), 616–624 (2023). 10.1038/s41586-023-06139-9.

[10] Menden, K. et al. Deep learning–based cell composition analysis from tissue expression profiles. Science Advances 6 (30), eaba2619 (2020). 10.1126/sciadv.aba2619.

[11] Aliee, H. & Theis, F. J. AutoGeneS: Automatic gene selection using multi-objective optimization for RNA-seq deconvolution. Cell Systems 12 (7), 706–715.e4 (2021). 10.1016/j.cels.2021.05.006.

[12] Chen, Y. et al. Deep autoencoder for interpretable tissue-adaptive deconvolution and cell-type-specific gene analysis. Nature Communications 13 (1), 6735 (2022). 10.1038/s41467-022-34550-9.

[13] Moffitt, J. R. et al. Molecular, spatial, and functional single-cell profiling of the hypothalamic preoptic region. Science 362 (6416), eaau5324 (2018). 10.1126/science.aau5324.

[14] Subramanian, A. et al. Gene set enrichment analysis: A knowledge-based approach for interpreting genome-wide expression profiles. Proceedings of the National Academy of Sciences 102 (43), 15545–15550 (2005). 10.1073/pnas.0506580102.

[15] Hwang, S. et al. HumanNet v2: Human gene networks for disease research. Nucleic Acids Research 47 (D1), D573–D580 (2019). 10.1093/nar/gky1126.

[16] Chen, J. et al. Transformer for one stop interpretable cell type annotation. Nature Communications 14 (1), 223 (2023). 10.1038/s41467-023-35923-4.

[17] Yang, F. et al. scBERT as a large-scale pretrained deep language model for cell type annotation of single-cell RNA-seq data. Nature Machine Intelligence 4 (10), 852–866 (2022). 10.1038/s42256-022-00534-z.

[18] Vaswani, A. et al. Guyon, I. et al. (eds) Attention is all you need. (eds Guyon, I.et al.) Advances in Neural Information Processing Systems, Vol. 30 (Curran Associates, Inc., 2017). URL https://proceedings.neurips.cc/paper files/paper/2017/file/3f5ee243547dee91fbd053c1c4a845aa-Paper.pdf.

[19] Devlin, J., Chang, M., Lee, K. & Toutanova, K. Burstein, J., Doran, C. & Solorio, T. (eds) BERT: pre-training of deep bidirectional transformers for language understanding. (eds Burstein, J., Doran, C. & Solorio, T.) (Association for Computational Linguistics, 2019). URL 10.18653/v1/n19-1423.

[20] Dosovitskiy, A. et al. An image is worth 16x16 words: Transformers for image recognition at scale. arXiv preprint 2010.11929 (2020). 10.48550/arXiv.2010.11929.

[21] Zhou, Y. et al. A foundation model for generalizable disease detection from retinal images. Nature 622 (7981), 156–163 (2023). 10.1038/s41586-023-06555-x.

[22] Cui, H. et al. scgpt: toward building a foundation model for single-cell multi-omics using generative ai. Nature Methods 1–11 (2024). 10.1038/s41592-024-02201-0.

[23] Lin, L. I.-K. A Concordance Correlation Coefficient to Evaluate Reproducibility. Biometrics? 45 (1), 255 (1989). 10.2307/2532051, 2532051.

[24] Safrastyan, A., Zu Siederdissen, C. H. & Wollny, D. Decoding cell-type contributions to the cfRNA transcriptomic landscape of liver cancer. Human Genomics 17 (1), 90 (2023). 10.1186/s40246-023-00537-w.

[25] Chen, S. et al. Cancer type classification using plasma cell-free RNAs derived from human and microbes. eLife 11, e75181 (2022). 10.7554/eLife.75181.

[26] Roskams-Hieter, B. et al. Plasma cell-free RNA profiling distinguishes cancers from pre-malignant conditions in solid and hematologic malignancies. npj Precision Oncology 6 (1), 28 (2022). 10.1038/s41698-022-00270-y.

[27] Tao, Y. et al. Cell-free multi-omics analysis reveals potential biomarkers in gastrointestinal cancer patients’ blood. Cell Reports Medicine 4 (11), 101281 (2023). 10.1016/j.xcrm.2023.101281.

[28] Humphries, A. & Wright, N. A. Colonic crypt organization and tumorigenesis. Nature Reviews Cancer 8 (6), 415–424 (2008). 10.1038/nrc2392.

[29] Barker, N. et al. Crypt stem cells as the cells-of-origin of intestinal cancer. Nature 457 (7229), 608–611 (2009). 10.1038/nature07602

[30] Bhowmick, N. A., Neilson, E. G. & Moses, H. L. Stromal fibroblasts in cancer initiation and progression. Nature 432 (7015), 332–337 (2004). 10.1038/nature03096.

[31] Boiarsky, R. et al. Single cell characterization of myeloma and its precursor conditions reveals transcriptional signatures of early tumorigenesis. Nature Communications 13 (1), 7040 (2022). 10.1038/s41467-022-33944-z.

[32] Wang, Y. et al. Plasma cell-free RNA characteristics in COVID-19 patients. Genome Research 32 (2), 228–241 (2022). 10.1101/gr.276175.121.

[33] Sun, J. et al. Early characterisation and prediction of liver diseases in pregnancy by plasma cell-free RNAs. Clinical and Translational Medicine 13 (10), e1439 (2023). 10.1002/ctm2.1439.

[34] Carsana, L. et al. Pulmonary post-mortem findings in a series of COVID-19 cases from northern Italy: A two-centre descriptive study. The Lancet Infectious Diseases 20 (10), 1135–1140 (2020). 10.1016/S1473-3099(20)30434-5.

[35] Rajamanickam, A. et al. Dynamic alterations in monocyte numbers, subset frequencies and activation markers in acute and convalescent COVID-19 individuals. Scientific Reports 11 (1), 20254 (2021). 10.1038/s41598-021-99705-y.

[36] Wilk, A. J. et al. A single-cell atlas of the peripheral immune response in patients with severe COVID-19. Nature Medicine 26 (7), 1070–1076 (2020). 10.1038/s41591-020-0944-y.

[37] Venet, F. et al. T cell response against SARS-CoV-2 persists after one year in patients surviving severe COVID-19. eBioMedicine 78, 103967 (2022). 10.1016/j.ebiom.2022.103967.

[38] Xu, G. et al. The differential immune responses to COVID-19 in peripheral and lung revealed by single-cell RNA sequencing. Cell Discovery 6 (1), 73 (2020). 10.1038/s41421-020-00225-2.

[39] Stephenson, E. et al. Single-cell multi-omics analysis of the immune response in COVID-19. Nature Medicine 27 (5), 904–916 (2021). 10.1038/s41591-021-01329-2.

[40] Sun, C., Sun, H., Zhang, C. & Tian, Z. NK cell receptor imbalance and NK cell dysfunction in HBV infection and hepatocellular carcinoma. Cellular & Molecular Immunology 12 (3), 292–302 (2015). 10.1038/cmi.2014.91.

[41] Cully, M. Reinvigorating exhausted T cells in hepatitis B infection. Nature Reviews Immunology 17 (4), 218–218 (2017). 10.1038/nri.2017.35.

[42] Ye, B. et al. T-cell exhaustion in chronic hepatitis B infection: Current knowledge and clinical significance. Cell Death & Disease 6 (3), e1694–e1694 (2015). 10.1038/cddis.2015.42.

[43] Moufarrej, M. N. et al. Early prediction of preeclampsia in pregnancy with cell-free RNA. Nature 602 (7898), 689–694 (2022). 10.1038/s41586-022-04410-z.

[44] Munchel, S. et al. Circulating transcripts in maternal blood reflect a molecular signature of early-onset preeclampsia. Science Translational Medicine 12 (550), eaaz0131 (2020). 10.1126/scitranslmed.aaz0131.

[45] Derisoud, E., Jiang, H., Zhao, A., Chavatte-Palmer, P. & Deng, Q. Revealing the molecular landscape of human placenta: a systematic review and meta-analysis of single-cell RNA sequencing studies. Human Reproduction Update dmae006 (2024). 10.1093/humupd/dmae006

[46] Castleman, J. S., Lip, G. Y. H. & Shantsila, E. Monocytes are increased in pregnancy after gestational hypertensive disease. Scientific Reports 12 (1), 10358 (2022). 10.1038/s41598-022-13606-2.

[47] Tang, Z. et al. CD36 mediates SARS-CoV-2-envelope-protein-induced platelet activation and thrombosis. Nature Communications 14 (1), 5077 (2023). 10.1038/s41467-023-40824-7.

[48] Nishikawa, M. et al. Massive image-based single-cell profiling reveals high levels of circulating platelet aggregates in patients with COVID-19. Nature Communications 12 (1), 7135 (2021). 10.1038/s41467-021-27378-2.

[49] Yang, L. et al. The signal pathways and treatment of cytokine storm in COVID-19. Signal Transduction and Targeted Therapy 6 (1), 255 (2021). 10.1038/s41392-021-00679-0.

[50] Klenk, C. et al. Platelet aggregates detected using quantitative phase imaging associate with COVID-19 severity. Communications Medicine 3 (1), 161 (2023). 10.1038/s43856-023-00395-6.

[51] Zhu, W. et al. Monoacylglycerol lipase promotes progression of hepatocellular carcinoma via NF-κB-mediated epithelial-mesenchymal transition. Journal of Hematology & Oncology 9 (1), 127 (2016). 10.1186/s13045-016-0361-3.

[52] Zhang, Z. et al. Serine catabolism generates liver NADPH and supports hepatic lipogenesis. Nature Metabolism 3 (12), 1608–1620 (2021). 10.1038/s42255-021-00487-4.

